# CRISPR-Cas12a targeting of ssDNA plays no detectable role in immunity

**DOI:** 10.1101/2022.03.10.483831

**Authors:** Nicole D. Marino, Rafael Pinilla-Redondo, Joseph Bondy-Denomy

## Abstract

CRISPR-Cas12a (Cpf1) is a bacterial RNA-guided nuclease that cuts double-stranded DNA (dsDNA) at sites specified by a CRISPR RNA (crRNA) guide. Additional activities have been ascribed to this enzyme *in vitro*: site-specific (*cis*) single-stranded DNA (ssDNA) cleavage and indiscriminate (*trans*) degradation of ssDNA, RNA, and dsDNA after activation by a complementary target. The ability of Cas12a to cleave nucleic acids indiscriminately has been harnessed for many applications, including diagnostics, but it remains unknown if it contributes to bacterial immunity. Here, we provide evidence that cleavage of ssDNA *in cis* or *in trans* by Cas12a is insufficient to impact immunity. Using LbCas12a expressed in either *Pseudomonas aeruginosa* or *Escherichia coli*, we observed that cleavage of dsDNA targets did not elicit cell death or dormancy, suggesting insignificant levels of collateral damage against host RNA or DNA. Canonical immunity against invasive dsDNA also had no impact on the replicative fitness of co-infecting ssDNA phage or plasmid *in trans*. Lastly, crRNAs complementary to invasive ssDNA did not provide protection, suggesting that ssDNA cleavage does not occur *in vivo* or is insignificant. Overall, these results suggest that CRISPR-Cas12a immunity predominantly occurs via canonical targeting of dsDNA, and that the other activities are *in vitro* irregularities.

## INTRODUCTION

CRISPR-Cas systems are prokaryotic adaptive immune systems that protect bacteria from mobile genetic elements (MGEs), such as phages and plasmids. Six distinct CRISPR-Cas types (I-VI) have been discovered, all featuring <u>C</u>RISPR-<u>a</u>ssociated (Cas) proteins and a CRISPR array that is established via acquisition of spacers from invasive nucleic acids (1). CRISPR arrays are transcribed and processed into CRISPR RNAs (crRNA) that associate with Cas proteins and direct them to their nucleic acid target (i.e. the protospacer) via complementary base pairing. In addition to sequence-specific targeting of DNA (Type I, II, V) (1) or RNA (Type II, III, and VI) (1, 2), some CRISPR-Cas systems have been ascribed a “collateral damage” phenotype that leads to the sequence-independent destruction of bystander ssRNA (Type III, VI) (3, 4) or DNA (Type III) (5). This indiscriminate cleavage can result in cell death (i.e. abortive infection) or dormancy (6), which prevents viral spread or escape. Bacterial nucleases involved in anti-phage immunity have also been shown to elicit abortive infection by targeting DNA and RNA indiscriminately, including Card1/Can2, Csm6, and NucC (7–10).

Cas12a (Type V-A CRISPR-Cas) is a programmable RNA-guided effector nuclease that creates dsDNA breaks within protospacers next to protospacer adjacent motifs (PAM) (11). Because of its high fidelity, distinct PAM requirements, and relatively short crRNA, Cas12a has been developed as an alternative to Cas9 for gene editing in bacteria, plants, and mammalian cells (12–14). Interestingly, Cas12a also cleaves ssDNA nonspecifically (i.e. *in trans*) upon binding to or cleaving its complementary target (i.e. *in cis*) *in vitro* (15–17). Recognition of the thymine-rich (TTTN) PAM in dsDNA initiates local strand separation and crRNA-target DNA hybridization (i.e. R-loop formation), which triggers conformational activation and exposure of the RuvC catalytic site (18–20). The RuvC domain then cleaves the non-target strand and target strand in rapid succession (19, 20). After cleavage, Cas12a releases the PAM-distal fragment of the target dsDNA but remains bound to the PAM-proximal fragment, which preserves the R-loop and catalytically active state (19–21). The exposed RuvC domain is free to cut any ssDNA that fits in its catalytic pocket, regardless of sequence. Notably, this *trans*-state can also be activated *in vitro* by binding complementary ssDNA in a PAM-independent manner (15, 16). Another study also reported degradation of ssRNA and dsDNA *in trans* after Cas12a activation, albeit at a slower rate than for ssDNA (17).

The remarkable ability of Cas12a to degrade ssDNA *in trans* upon activation has led to its rapid development in diagnostics as a DNA endonuclease-targeted CRISPR trans reporter (DETECTR) (15); however, recent work *in vitro* suggests its kinetics were overestimated (22). Nonetheless, the success of these applications and a recent explosion in the functional and evolutionary diversity of the Cas12 (Type V) superfamily have led to a flurry of new detection tools with similar capabilities (23–28). Although the collateral damage observed *in vitro* for these nucleases is well-suited for diagnostics, it raises some concerns about their use in gene therapy (29).

Cas12a *cis*- and *trans*-cleavage *in vitro* has been carefully elucidated using structural and biochemical studies (18–20, 30), however no significance of this activity has been described during cellular immunity in bacteria. Bacteria encounter ssDNA in many forms, including bacterial or MGE replication forks and transcription bubbles, ssDNA phages, and dsDNA conjugative elements that transfer through a ssDNA intermediate. Indiscriminate ssDNA cleavage could therefore yield many outcomes: i) destruction of ssDNA phage or conjugative elements, ii) targeting of dsDNA phage and plasmid during replication or transcription, or iii) self-targeting during bacterial replication or transcription, possibly resulting in cell dormancy or death. Indiscriminate cleavage of host RNA or dsDNA upon activation could likewise confer immunity through abortive infection. Here, we provide evidence that Cas12a does not exhibit indiscriminate cleavage activities that impact bacterial immunity. To address these questions, we programmed LbCas12a to target dsDNA phage, ssDNA phage, or conjugated plasmids in *E. coli* or *P. aeruginosa*. Although we observed strong canonical immunity against dsDNA phage and plasmid, this did not elicit abortive infection or impact the replicative fitness of co-infecting ssDNA phage or plasmid *in trans*. crRNAs complementary to invasive ssDNA also did not provide direct protection. Altogether, these results suggest that cleavage of ssDNA, ssRNA, and host dsDNA does not detectably contribute to immunity in the cell.

## MATERIALS AND METHODS

### Bacterial strains, plasmids, phages and media

The bacterial strains, plasmids and phages used in this study are described in Supplemental Table S1. Bacterial strains were cultured on lysogeny broth (LB) agar or liquid media at 37°C or 30°C as indicated. For *Pseudomonas aeruginosa* PAO1 strain, extrachromosomal plasmids were maintained using antibiotics at the indicated concentrations: pHERD30T and derivatives, 50 μg/ml gentamicin; pKJK5 (pTrans), 60 μg/ml trimethoprim. For *E. coli* strains, extrachromosomal plasmids were maintained as follows: pHERD20T, 100 μg/ml carbenicillin (strain BW25113); pHERD30T and derivatives, 15 μg/ml gentamicin, pKJK5 (pTrans), 50 μg/ml kanamycin (strains S17-1 and CSH26). The following strains were grown in media supplemented with antibiotics: *E. coli* S17-1: 30 μg/ml streptomycin; *E. coli* CSH26: 30 100 μg/ml rifampicin, 100 μg/ml nalidixic acid; PAO1 tn7::LbCas12a attB::crRNA24, 50 μg/ml tetracycline. Inducers were added to the media at the following concentrations: 1 mM IPTG and 0.3% arabinose unless otherwise indicated. M13 phage was purchased from ATCC. λ_vir_ was a gift from Ry Young, and Pf4 phage was a gift from Paul Bollyky.

### Construction of plasmids and strains

To generate the PAO1 tn7::LbCas12a strain (Supplementary Figure S1), LbCas12a was first amplified from pMBP-LbCas12a (Addgene #113431) and cloned into pUC18-mini-Tn7T-lac. pUC18-Tn7T-LbCas12a was co-electroporated with pTNS3 plasmid into the *P. aeruginosa* PAO1 strain according to previously published protocols (31, 32). The resulting transformants were plated on LB agar supplemented with 30 μg/ml gentamicin to select for integration of the pUC18-Tn7T-LbCas12a vector into the bacterial chromosome. Colonies were screened for integration at the Tn7 site and cured of the gentamicin-resistance cassette using Flp-mediated marker excision, as previously described (31). Clones sensitive to gentamicin were then passed serially in LB to cure them of pFLP2 and confer carbenicillin sensitivity.

crRNAs complementary to DMS3 and JBD30 phages or RFP control were cloned into the HindIII and NcoI sites of pHERD30T. For experiments in which plasmid-borne crRNAs were used, the resulting plasmids were introduced into PAO1 tn7::LbCas12a directly via electroporation and selection on LB supplemented with 50 μg/ml gentamicin. To integrate crRNAs into the chromosome, the region encompassing the *araC* gene, pBAD promoter, and crRNA was amplified from pHERD30T-crRNA vectors and cloned into the mini-CTX2 vector. The resulting plasmids were electroporated into the PAO1 tn7::LbCas12a strain, and integration into the chromosome was selected for using 15 μg/ml tetracycline according to established protocols (33).

To generate the *E. coli* BW25113 F’ strain, BW25113 was first mated with TOP10F *E. coli* and plated on M9 minimal media supplemented with 10 μg/ml tetracycline to select for retention of F’ in BW25113 and death of the TOP10F leu-autotroph. LbCas12a was amplified from pMBP-LbCas12a and cloned into pTN7C185. The resulting pTN7-LbCas12a vector and pTNS3 were introduced into BW25113 F’ via triparental mating, and integration of pTN7-LbCas12a into the BW25113 F’ chromosome was selected by plating cells on LB supplemented with 10 μg/ml gentamicin (Supplementary Figure S1). Colonies were screened for vector integration into the Tn7 site. crRNAs complementary to λ_vir_ or M13 were cloned into the NcoI and HindIII sites of pHERD20T and introduced into BW25113 F’ via chemical transformation.

### Liquid culture phage infections

Overnight cultures of *P. aeruginosa* were diluted 1:100 in LB supplemented with 10 mM MgSO4, 1 mM IPTG, 0.3% arabinose and 50 μg/ml gentamicin. 90 μl of diluted bacteria were then infected with 10 μl DMS3 phage diluted ten-fold in SM phage buffer in a 96-well Costar plate. These infections proceeded for 24 hours in a Synergy H1 microplate reader with continuous, double orbital shaking at 37°C. PAO1 tn7::LbCas12a pHERD30T-crRNA-RFP was used as the non-targeting control and PAO1 tn7::LbCas12a pHERD30T-crRNA-DMS3 was used for targeting. Dilutions of bacteria and phage were plated to determine the number of colony forming units (CFUs) and plaque forming units (PFUs), respectively. The multiplicity of infection was determined by dividing the number of PFUs by CFUs.

### Generation of phage escapers

High titer preparations of JBD30 phage (which encodes the same protospacer present in DMS3) were mixed with overnight cultures of PAO1 tn7::LbCas12a attB::crRNA-DMS3 in top agar and plated on LB supplemented with 1 mM IPTG and 0.3% arabinose. Cultures were grown overnight at 30°C, and resulting plaques were isolated with a sterile pipette tip and resuspended in SM phage buffer with chloroform. The resulting phages were plaqued twice on targeting media and tested for plaque formation on targeting and non-targeting strains. The protospacer regions of *bona fide* phage escapers and wildtype phage were Sanger sequenced and aligned using EMBOSS Needle.

### Phage plaque assays

Bacterial lawns were generated by adding 150 μl bacteria from an overnight culture to 4 mls of 0.7% top agar and spreading the mixture on LB agar supplemented with 10 mM MgSO_4_ (*P. aeruginosa*) or 10 mM MgSO_4_ and 5 mM CaCl_2_ (*E. coli*). Phages were diluted in SM phage buffer (JBD30 and DMS3) or LB supplemented with 10 mM MgSO_4_ and 5 mM CaCl_2_ (λ_vir_ and M13). 3 μl of phage was spotted onto bacterial lawns and grown at 30°C or 37°C overnight, respectively. The identity and protospacer sequences of phage stocks were confirmed by PCR and Sanger sequencing.

### Plasmid conjugation assay

Plasmid conjugation assays were carried out as solid surface filter matings. Five different *E. coli* S17-1 plasmid-mobiliser strains (donors) were independently mated with the two recipient strains in triplicate. Wild type PAO1 (wt) or PAO1 tn7::LbCas12a attB::crRNA-DMS3 was used as the recipient strain. Each donor strain carried unmodified pHERD30T or a derivative encoding the complementary protospacer sequence on the leading strand (LDS) or lagging strand (LGS) with or without a PAM (Supplementary Figure S2). The strains were grown overnight in LB broth supplemented with the appropriate antibiotics and CRISPR-Cas inducers. Donor and recipient cell suspensions (500 μL) were washed three times, mixed at a 1:1 ratio and spun down. The pellet was then resuspended in 100 μL LB and transferred onto sterile 0.2 μm nitrocellulose filters (Avantec) placed over nonselective LB agar plates with inducers. The bacterial mating mixtures were incubated for 3h at 37°C to allow conjugation. Filters were then washed with 2mL PBS to recover cells. Serial dilutions were subsequently plated on selective LB-agar plates with appropriate antibiotics (and no inducers) to discriminate between donor, recipient and transconjugant colony forming units (CFUs). The frequency of plasmid transfer was calculated by dividing the number of transconjugant CFUs by the number of donor CFUs.

### Plasmid co-conjugation assay

The co-conjugation experiments were performed as solid surface filter matings, in triplicate, using *E. coli* strains S17-1 and CSH26 as donors and PAO1 tn7::LbCas12a attB::crRNA-DMS3 (targeting) as the recipient. The pTrans plasmid was simultaneously conjugated with pTarget (which encodes a complementary protospacer) or pTarget(-PS) (which lacks a complementary protospacer), into the recipient strains. Donor and recipient strains were grown overnight using the appropriate selection and CRISPR-Cas induction (recipient only). The donor and recipient cell suspensions (500 μL) were washed 3 times, mixed at a 1:1:1 ratio and spun down. The pellet was then resuspended in 100 μL LB and transferred onto sterile 0.2 μm nitrocellulose filters (Avantec) placed over nonselective LB agar plates with inducers and no antibiotic selection. The bacterial mating mixtures were incubated for 3h at 37°C to allow conjugation. Filters were then washed with 2mL PBS to recover cells. Serial dilutions were subsequently plated on selective LB-agar plates with appropriate antibiotics to discriminate between donor, recipient and transconjugant CFUs. The frequency of plasmid transfer was calculated by dividing the number of transconjugant CFUs by the number of donor CFUs.

### In vitro transcription

crRNAs were synthesized for *in vitro* cleavage experiments using the HiScribe T7 Quick High Yield RNA Synthesis Kit (New England Biolabs; NEB). Templates for *in vitro* transcription were generated by hybridizing a ssDNA oligo with the T7 promoter to another oligo with the complementary T7 promoter sequence fused to the LbCas12a repeat and spacer in annealing buffer. Approximately 1 μg template was added to the HiScribe reaction and incubated at 37°C for 16-18 hours. Reactions were purified using the Qiagen miRNeasy kit and resulting concentrations were measured using Qubit RNA Broad Range Assay Kit (Thermo Fisher) and the Qubit Fluorometer (Invitrogen). Purified RNA was loaded onto TBE-urea gels (BioRad) and resolved by electrophoresis in TBE buffer with low range ssRNA ladder (NEB). Gels were post-stained with SYBR gold for 1 hr at room temperature and imaged using BioRad Gel Doc EZ Imager.

### Pf4 propagation and purification

LB broth (3 mL) was inoculated with a colony of PAO1 and grown for 4h, 37°C at 250 revolutions per minute (rpm). 3 μL Pf4 phage stock was used to infect each culture (except for one uninfected control). Infected cultures were grown for an additional 1-2 hrs and transferred to a flask with 75 mL LB broth for 24-48 hours, 37°C at 250 rpm. Cells were harvested by centrifugation at 8,000 x g for 10 min, 4°C. The supernatant was collected and treated with 1 μg/mL DNase I for 2 hrs, 37°C. Cells were centrifuged again at 8,000 x g for 10 min, 4°C and filter sterilized with a 0.22 μm filter. The filtrate was supplemented with 5M NaCl and incubated at 4°C for 4 hrs. Polyethylene glycol (PEG 8000) was added at a final concentration of 4% (w/v), and the sample was left gently rotating at 4°C overnight. To pellet phage, the samples were centrifuged at 13,000 x g for 20 min at 4°C. Pellets were resuspended in 30 mls 10mM Tris 1mM EDTA, pH 8.0 (TE) buffer and centrifuged again at 15,000 x g at 4°C. The supernatant was collected and supplemented with a quarter-volume of 20% PEG 8000 and 2.5 M NaCl and left overnight at 4°C. Phage was harvested by centrifuging at 15,000 x g, 4°C, 20 min. The resulting pellets were resuspended in 2 mls PBS and dialyzed overnight at 4°C in 1L PBS using SnakeSkin 10,000 MWCO dialysis tubing (Thermo Fisher). Buffer was refreshed and samples were dialyzed again overnight. Virion viability and yield was determined by plaque assay on PAO1.

### Genomic DNA extraction

DNA from purified Pf4 virions was extracted by mixing phage preparations 1:1 in lysis solution (0.4% sodium dodecyl sulfate, 400 μg/ml Proteinase K, 200 μg/ml RNase A) and incubating at 37°C for 1 hr. Samples were then incubated at 50°C for 30 minutes, supplemented with 0.2M NaCl, and subjected to a standard phenol-chloroform extraction. PAO1 genomic DNA was extracted from saturated cultures using the Qiagen DNeasy Ultraclean Microbial Kit according to the manufacturer’s instructions.

### In vitro cleavage experiments

*In vitro* Cas12a cleavage assays were performed using EnGen Lba Cas12a (NEB #M0653T) and crRNAs that were synthesized commercially (IDT) or with the HiScribe T7 Quick High Yield RNA Synthesis Kit (NEB). dsDNA templates were amplified from plasmids and purified using DNA Clean and Concentrator Kit (Zymo). 30 nM EnGen LbCas12a was pre-incubated with 30 nM crRNA in NEBuffer 2.1 for 10 minutes at 25°C. Substrate DNA (i.e. dsDNA template, M13mp18 ssDNA (NEB #N4040S) or purified Pf4 DNA) was added and incubated at 37°C for 2 hours. 1 μl Proteinase K (50 mg/ml) was added and the reaction was incubated at room temperature for 10 minutes. *In vitro* cleavage experiments using shrimp dsDNAse (NEB; EN0771), EcoRI-HF (NEB; R3101S), and Exonuclease I (NEB; M0293S) were performed for 2 hours at 37°C and heat inactivated at 65°C. Reactions were loaded onto 1% TAE agarose gels and resolved by electrophoresis. Gels were post-stained with SYBR Gold (Invitrogen) for 1 hr at room temperature and imaged using BioRad Gel Doc EZ Imager.

## RESULTS

### Cas12a targeting does not elicit abortive infection or prevent escaper phage emergence

Diverse orthologs of Cas12a have been shown to *cis*- and *trans*-cleave ssDNA *in vitro*, including LbCas12a (*<u>L</u>achnospiraceae <u>b</u>acterium ND 2006*), AsCas12a (*<u>A</u>cidaminococcus* <u>s</u>p. *BV3L6*) and FnCas12a (*<u>F</u>rancisella <u>n</u>ovicida*) (15–17). The bacteria in which these orthologs are naturally found are either genetically intractable or lack known plasmids and phages that can be manipulated to assess *cis*- and *trans*-cleavage. Therefore, we engineered the *P. aeruginosa* lab strain PAO1 to express IPTG-inducible LbCas12a from the chromosome and arabinose-inducible crRNAs programmed to target phage (Supplementary Figure 1). *P. aeruginosa* has previously been used as a host for MbCas12a (*Moraxella bovoculi*) to target phage and identify anti-CRISPR proteins (34).

To determine if Cas12a triggers abortive infection upon cleaving its complementary target, we first infected targeting and non-targeting strains of *P. aeruginosa* with phage DMS3 at varying multiplicities of infection (MOI) and assessed bacterial growth over time. If phage targeting by Cas12a triggers abortive infection by *trans*-cleaving host nucleic acids, infecting cells with a high MOI (in which most cells are infected) would cause population collapse before phage replication is complete. We observed no significant deviation in the growth kinetics or number of colony forming units (cfus) during this experiment (Figure 1A). Notably, Cas12a immunity was incredibly robust, protecting cells across a 10^7^-fold MOI range. Due to the strength of immunity and the absence of cell death, phage escapers emerged from high MOI infections (Figure 1B). These escaper phages had single point mutations in the +2 or +3 position of the seed (PAM-proximal) region of the protospacer, which is consistent with the high sequence fidelity required in this region for successful targeting. Altogether, these data show that LbCas12a targeting does not induce abortive infection or prevent the emergence of escapers.

**Figure 1.**
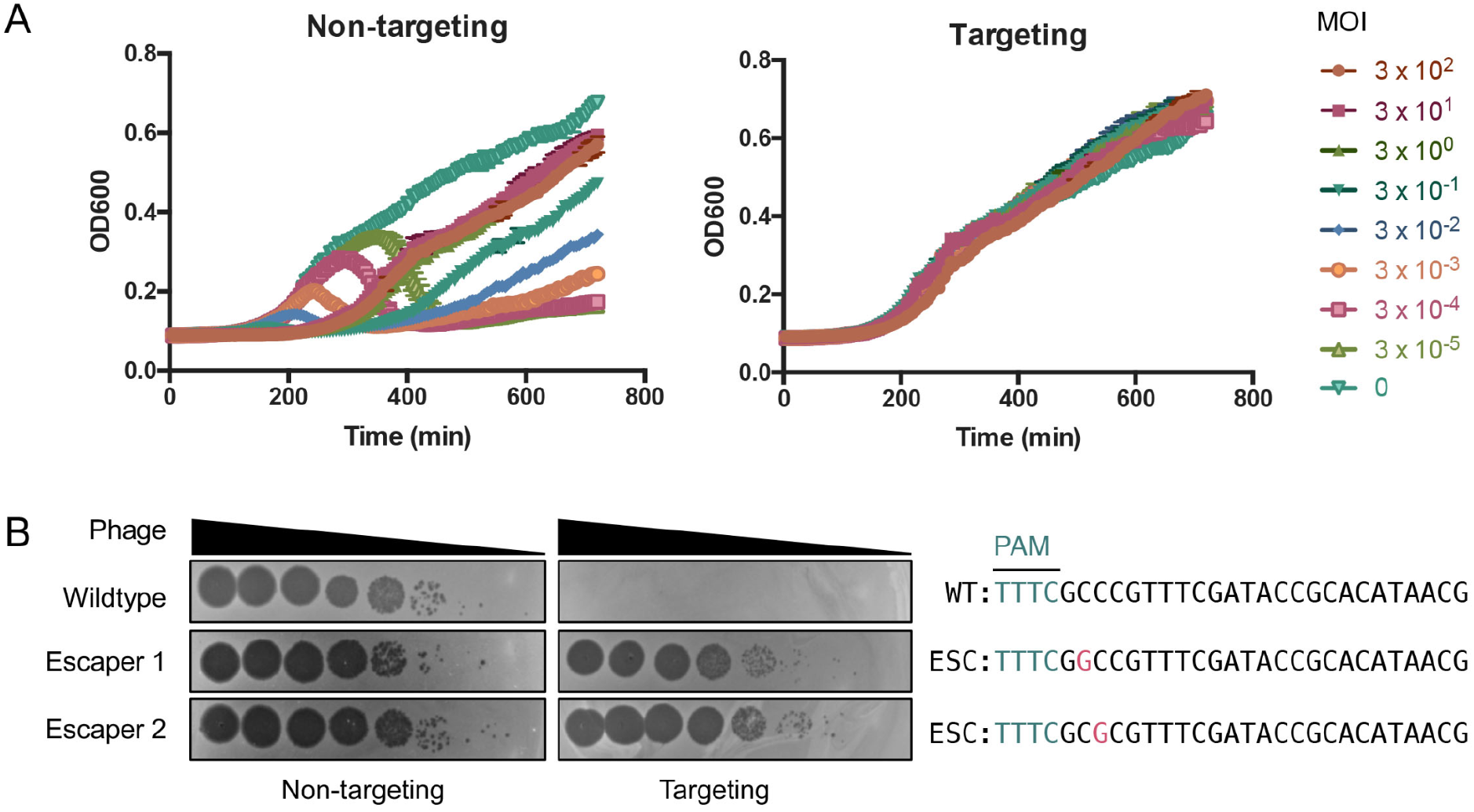
Cas12a targeting does not elicit abortive infection or prevent the rise of phage esapers. (A) Growth curves of *P. aeruginosa* PAO1 tn7::LbCas12a strain expressing targeting or non-targeting crRNAs infected with ϕDMS3 at multiplicities of infection (MOI) indicated in legend. (B) Phage plaque assays with ten-fold serial dilutions of wildtype or mutated ϕ JBD30 phage. Bacterial clearance (black) indicates phage replication. Sequences of the targeted phage gene show mutated nucleotides (magenta) in phages able to escape Cas12a targeting. PAM, protospacer adjacent motif (teal); WT, wildtype phage; ESC, escaper phage.

### Cas12a does not cis-cleave invasive ssDNA in cells

We next tested the ability of Cas12a to cleave ssDNA in cells through direct complementary base pairing. We exposed cells to plasmids and ssDNA phages, which enter and exit the bacterium as ssDNA but replicate via a dsDNA intermediate (35). First, we performed conjugation assays using plasmids with the complementary protospacer on the leading or lagging strand. Only the leading strand transfers as ssDNA from the donor to the recipient cell during conjugation; therefore, only plasmids with the complementary sequence on the leading strand should be susceptible to Cas12a ssDNA cleavage. Because Cas12a requires the PAM for dsDNA unwinding but not conformational activation, we used target constructs with or without the PAM to discriminate between cleavage of dsDNA and ssDNA, respectively (Figure 2A and Supplementary Figure S2). As expected, LbCas12a targeted conjugated plasmid with PAM(+) protospacers regardless of the orientation, which is consistent with cleavage of the dsDNA form (Figure 2A and Supplementary Figure S3). However, we did not detect any targeting of conjugated plasmids with PAM(-) protospacers present on the leading or lagging strand, suggesting the ssDNA strand was not significantly cleaved upon entering the cell.

**Figure 2.**
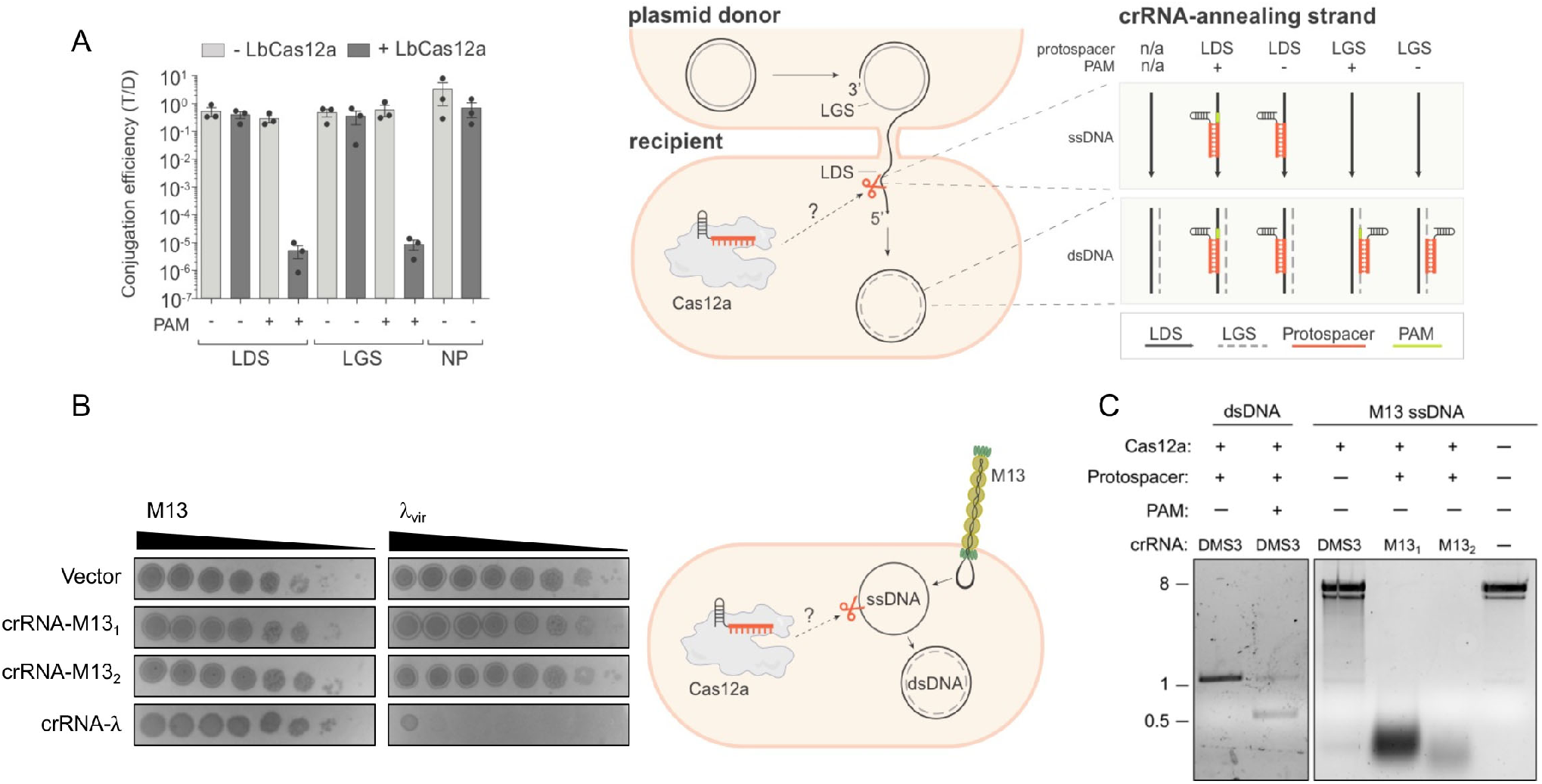
Cas12a does not detectably cis-cleave invasive ssDNA in bacteria. (A) Conjugation efficiency for plasmid cis-targeting in *P. aeruginosa* PAO1 strain. During conjugation, the donor is nicked and the leading strand (LDS) is transferred as ssDNA to the wildtype (-LbCas12a) or LbCas12a-expressing (+LbCas12a) recipient cell via the mating pilus. LbCas12a was co-expressed with crRNAs complementary to a DMS3 protospacer cloned into the leading (LDS) or lagging (LGS) strand, with (+) or without (-) the correct protospacer adjacent motif (PAM). T/D indicates the ratio of transconjugants to donors. Data are presented as mean±SEM (n = 3). LDS, complementary protospacer is on leading strand; LGS, complementary protospacer is on lagging strand; NP, no protospacer on plasmid. (B) Plaque assay for phage cis-targeting in *E. coli* BW25113 F’ strain. Bacterial lawns were infected with dsDNA λ_vir_ or with M13, which enters and leaves the cell as ssDNA but replicates via a dsDNA intermediate. LbCas12a was co-expressed in bacteria with crRNA complementary to M13 positive strand (without PAM) or λ_vir_ (with PAM) during phage infection. Bacterial clearance (black) indicates phage replication. (C) *In vitro* LbCas12a cleavage assay on dsDNA template encoding DMS3 protospacer or M13 ssDNA.

We then sought to perform ssDNA cleavage experiments in *P. aeruginosa* using the filamentous phage Pf4, which similarly enters and leaves the cell as linear ssDNA and replicates as a dsDNA intermediate (36, 37). We infected our model strain with varying titers of Pf4 and again observed targeting only when using crRNAs corresponding to PAM(+) (i.e. dsDNA) protospacers (Supplementary Figure S4). However, we found that DNA purified from Pf4 virions was surprisingly recalcitrant to *trans*-cleavage *in vitro* (Supplementary Figure S4D). We attempted to confirm the single-stranded nature of the phage DNA using a panel of nucleases, with PAO1 genomic DNA included for comparison. Surprisingly, we found that DNA purified from Pf4 was susceptible to dsDNAse but not exonuclease I, which cleaves linear ssDNA nonspecifically (Supplementary Figure S4E and S4F). Although DNA purified from Pf4 resolved around the expected size (12.4 kb) (38) and was smaller than PAO1 genomic DNA, its susceptibility to EcoRI and dsDNAse suggests that the genome harvested is dsDNA. As a result, we could not definitively conclude the nature of this DNA or the suitability of this phage for assessing ssDNA cleavage.

We therefore established a second bacterial model for LbCas12a in the *E. coli* strain BW25113 F’ and used dsDNA phage λ_vir_ and ssDNA phage M13 to assess cleavage activities. M13, like Pf4, is a filamentous phage that enters and exits the bacterium as ssDNA but replicates as dsDNA via rolling-circle replication (39, 40). M13 ssDNA was demonstrated to be rapidly degraded *in cis* and *in trans* by LbCas12a *in vitro* (15–17) and M13 has been used as a model for Type I-E CRISPR-Cas acquisition and targeting in *E. coli* (41). As with our *P. aeruginosa* model, we engineered *E. coli* to express IPTG-inducible LbCas12a from the chromosome and arabinose-inducible crRNAs (Supplementary Figure S1). When we expressed crRNAs complementary to PAM(+) protospacers in λ_vir_, we saw a 10^7^-fold reduction in phage titer (Fig 2B). When we expressed crRNAs complementary to PAM(-) protospacers in M13, however, we saw no reduction in plaque formation, even though these same crRNAs enabled cleavage of M13 ssDNA *in vitro* (Figures 2B, 2C). These results indicate that LbCas12a readily targets dsDNA, but not ssDNA phage or plasmids in cells, suggesting that the robust degradation of M13 ssDNA observed *in vitro* does not recapitulate during bacterial infection.

### Cas12a does not trans-cleave invasive ssDNA in cells

Although we could not detect *cis* ssDNA cleavage, we next probed for *trans* ssDNA cleavage that could result after cleavage of a dsDNA target. To this end, we simultaneously conjugated two plasmids—pTarget, which has the correct protospacer and PAM, and pTrans, which has no protospacer—into the same recipient cells and selected for transconjugants using antibiotic selection specific to each plasmid. As a control, we used pTarget(-PS), which is identical to pTarget but lacks the protospacer and PAM. If cleavage of plasmid with the correct target elicits indiscriminate immunity against ssDNA, the number of Cas12a-expressing transconjugants retaining pTrans should be lower during co-conjugation with pTarget than for the pTarget(-PS) control. While conjugation of pTarget was decreased by >3 orders of magnitude relative to pTarget(-PS), this did not impact pTrans acquisition (Fig 3A, Supplementary Figure S5). We then tested for *trans*-cleavage of ssDNA phage by singly infecting or co-infecting *E. coli* with dsDNA phage λ_vir_ and ssDNA phage M13. When we expressed LbCas12a and λ_vir_-targeting crRNA during infection, we observed strong reduction in λ_vir_ titer but no effect on the plaque forming ability of a co-infecting M13 phage, even across a 10^7^-fold range of phage titers (Figure 3B, Supplementary Fig S6). Altogether, we conclude that the potential for *trans* cleavage to contribute is undetectable.

**Figure 3.**
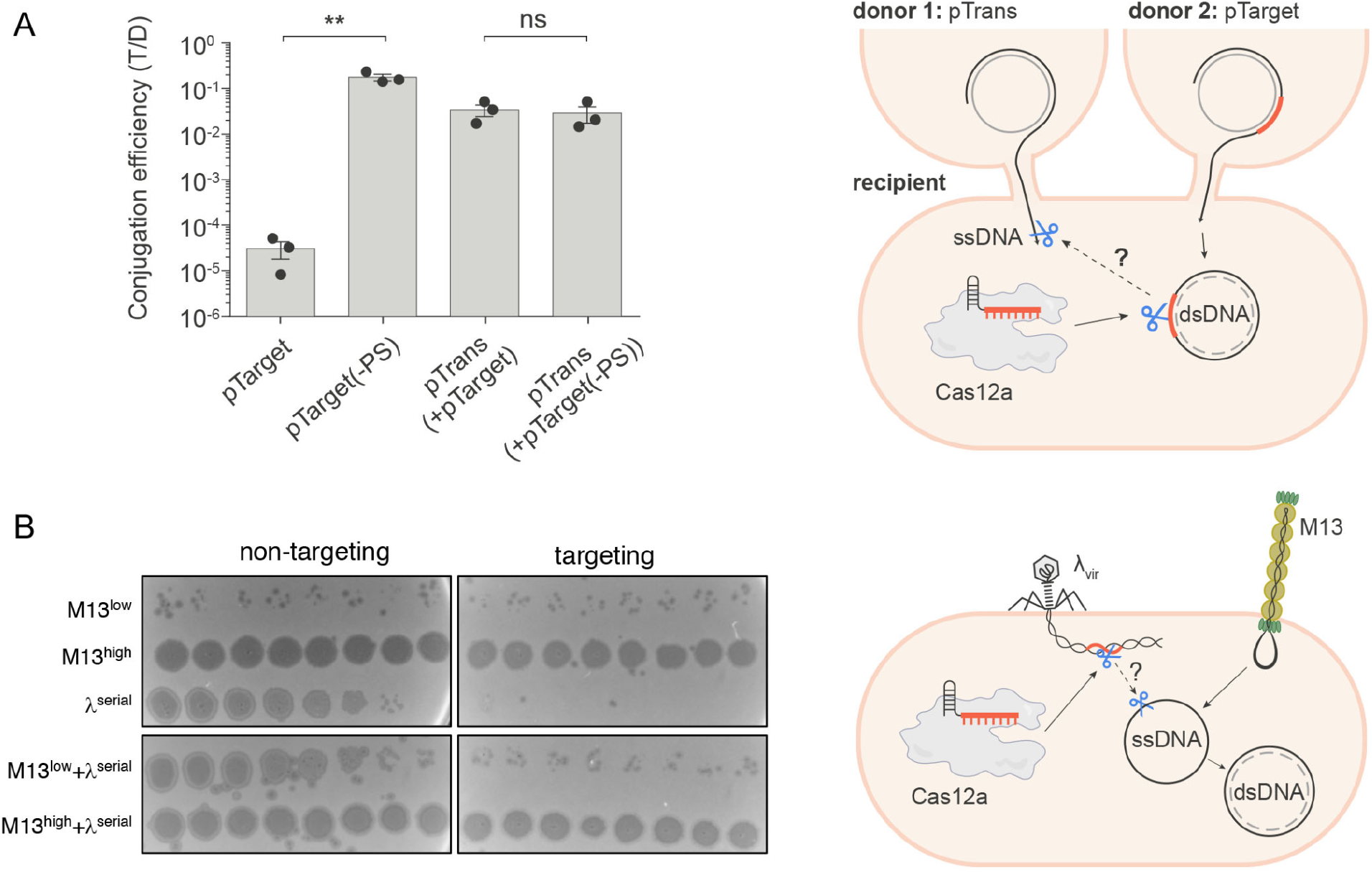
Cas12a does not trans-cleave invasive ssDNA in bacteria. (A) Right: Schematic for plasmid co-conjugation and cleavage in *P. aeruginosa* PAO1 strain. Two plasmids were conjugated simultaneously from different donors into the same recipient cell. LbCas12a was co-expressed in the recipient cell with crRNA complementary to pTarget but not pTrans. A parallel co-conjugation mating using pTrans and pTarget(-PS) (identical to pTarget but lacking the protospacer and PAM) was carried out as a control. Conjugation efficiency was assessed using antibiotic selection specific to each plasmid and represented as the transconjugant to donor ratio (T/D). Data are presented as mean ± SEM (*n* = 3); \*\**p*=0.0036 by unpaired, two-sided Student’s *t* test; ns, not significant (*p*>0.05) (B) Right: Schematic for phage co-infection in *E. coli.* Bacterial lawns of *E. coli* BW25113 F’ strain co-expressing LbCas12a and λ-targeting or non-targeting crRNA were singly infected or co-infected with ssDNA phage M13 and dsDNA phage λvir. Strains were infected with either low titer (low), high titer (high), or ten-fold serial dilutions (serial) of phage as indicated.

## DISCUSSION

Although Cas12a is generally used to create staggered dsDNA breaks at specified sites, indiscriminate DNA and RNA shredding *in vitro* has been observed after target cleavage (15–17). This non-specific *trans*-cleavage has been repurposed for diagnostics, but it is not known if it occurs in cells or contributes to bacterial immunity. Here, we provide evidence that ssDNA *cis*- or *trans*-cleavage by Cas12a is insufficient to contribute to immunity against ssDNA plasmid or phage. The reasons for this are unclear, but it may be due to protection of ssDNA by DNA-binding proteins that exclude Cas12a or the rapid conversion of invasive ssDNA to dsDNA replicative forms in the cell (40, 42–45). DNA or RNA polymerase may likewise displace “activated” Cas12a protein from its target. Consistent with this, ssDNA cleavage rates are not as high as initially reported and may be insignificant given the transient nature of naked ssDNA in the cell (22). We also saw no evidence of LbCas12a triggering cell dormancy or death, suggesting it does not significantly target host dsDNA or RNA. Given that the degree of Cas12a trans-cleavage varies across buffer conditions (17), natural conditions in the cell may also maintain LbCas12a in a conformational state that is less prone to *trans*-cleavage.

The results shown here are consistent with previous reports showing high specificity and relatively infrequent off-target effects for Cas12a in cells (46–49). These findings suggest that indiscriminate Cas12a cleavage activities might be *in vitro* artefacts, perhaps due to the absence of cellular factors like ssDNA binding proteins and polymerases. It must be noted, however, that the strains used here are not natural hosts for Type V-A CRISPR systems. Because native hosts for Cas12a are either genetically intractable or lack tractable plasmids and phages, we opted to express inducible LbCas12a in *E. coli* and *P. aeruginosa* and test for ssDNA cleavage using plasmids and phages that infect these species. Although we did not observe *cis*- or *trans*-ssDNA cleavage across a range of conditions, it’s possible that Cas12a ssDNAse activity exists only in its native hosts or against certain species of ssDNA.

Nonetheless, the rapid degradation of M13 ssDNA by Cas12a observed *in vitro* does not recapitulate *in vivo*, showing that CRISPR-Cas activities in cells cannot be understood through *in vitro* studies alone. Although indiscriminate cleavage activities *in vitro* have raised concerns about the use of Cas12a in gene editing, Cas12a has been generally regarded as an accurate enzyme in mammalian cells (50) and our results suggest the absence of rampant ssDNA cleavage during anti-phage/plasmid immunity. Cas12a therefore remains a powerful and versatile tool for the myriad of applications that already exist and are still to come.

## DATA AVAILABILITY

All data and relevant information will be made available upon request.

## ACKNOWLEDGMENTS

We thank members of the Bondy-Denomy lab for thoughtful discussions, including Sukrit Silas for experimental advice. We thank Jason M. Peters for the pTN7C185 vector and advice on strain engineering. We also thank Paul Bollyky, Joliene Sweere, Pat Secor and Alison Coluccio for their help with Pf4 phage. Author contributions: N.D.M. conceived of the project, engineered the bacterial strains and performed phage infection and *in vitro* experiments. R.P.-R. performed all conjugation experiments and made the figure schematics. N.D.M., R.P.-R. and J.B.-D. all contributed to experimental design and writing of the manuscript. J.B.-D. supervised the project.

## FUNDING

N.D.M. was supported by the National Institutes of Health F32 [F32GM133127]. R.P.-R. was supported by the Lundbeck Foundation (Lundbeckfonden) postdoctoral grant [R347-2020-2346]. J.B.-D. was supported by a National Institutes of Health Director’s Early Independence Award [DP5-OD021344, R01GM127489].

## CONFLICTS OF INTEREST

J.B.-D. is a scientific advisory board member of SNIPR Biome and Excision Biotherapeutics and a scientific advisory board member and co-founder of Acrigen Biosciences. The Bondy-Denomy lab receives research support from Felix Biotechnology. J.B.-D. and N.D.M. have filed patents on technology related to Cas12a anti-CRISPR proteins.

