## Supplementary Materials for "CRISPR-Cas12a targeting of ssDNA plays no detectable role in immunity"

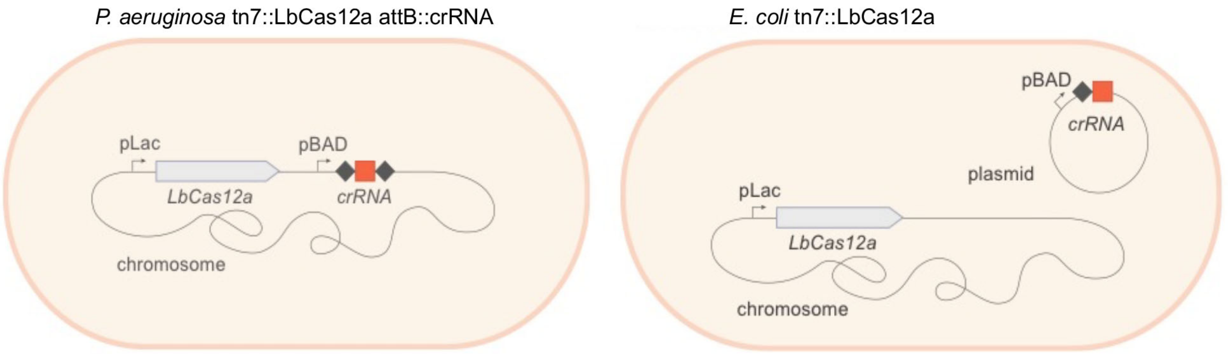


Figure S1. Schematic of the strains used in this study. *Pseudomonas aeruginosa* PAO1 strain was engineered to express chromosomally integrated LbCas12a and crRNA. *E. coli* BW25113 strain was engineered to express chromosomally integrated LbCas12a and plasmid-borne crRNA.

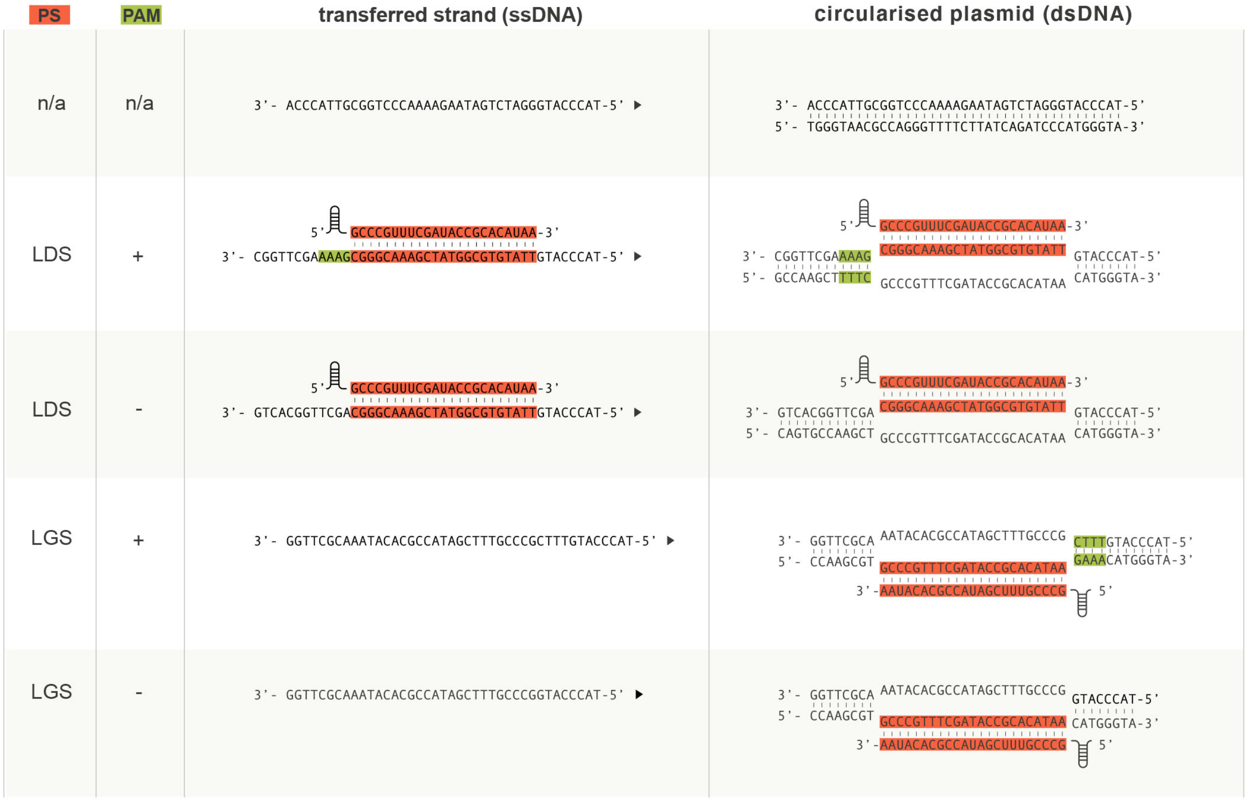


Figure S2. Schematic for crRNA and protospacers used for conjugation experiments. Five different pHERD30T-based plasmid constructs were used to assess for ssDNA cis-cleavage during conjugation. The spacer-protospacer matches are shown in red and the PAM sequence in green. The left panel depicts the presence or absence of complementary base-pairing between the crRNA and protospacers in the leading strand (transferred strand); the black arrow head indicates the direction of transfer during conjugation from the donor to the recipient cell. The right panel illustrates base pairing between the crRNA and protospacers in the plasmids once they become dsDNA. LDS, complementary protospacer is on the leading strand; LGS, complementary protospacer is on the lagging strand; PS, protospacer; PAM, protospacer adjacent motif.

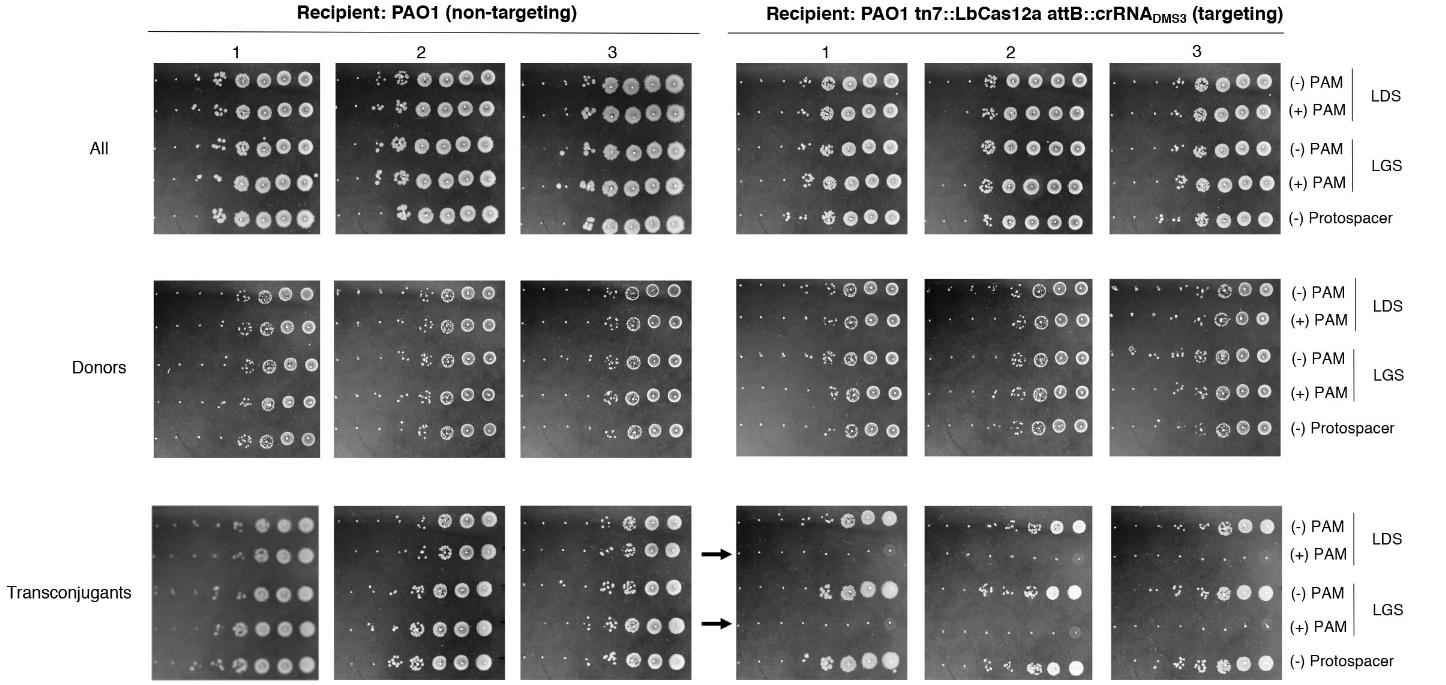


Figure S3. LbCas12a does not cis-cleave ssDNA during plasmid conjugation. Conjugation assays for ssDNA *cis*-cleavage. Colony forming units (10-fold dilution series) of transconjugants (bottom), donors (middle) and all (top) cells were grown in the appropriate selective media after the conjugative filter mating. Numbered columns indicate the three replicates for the combination of matings performed with either the non-targeting recipient (left) or the targeting recipient (right). Dilution rows start at $10^{-2}$ for “all” and $10^{0}$ for “donors" and “transconjugants”. LDS, the complementary protospacer is on the leading strand; LGS, the complementary protospacer is on the lagging strand; (+/-) PAM indicates the presence/absence of the correct protospacer adjacent motif (PAM). Black arrows indicate where targeting is observed.

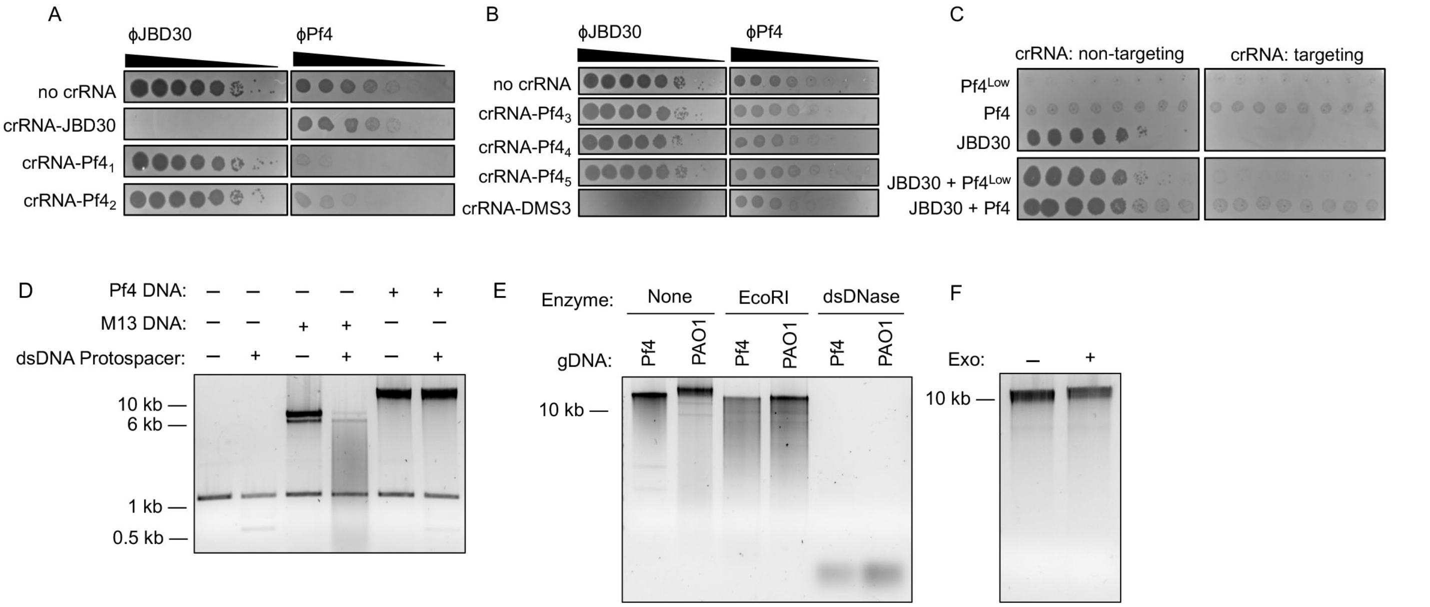


Figure S4. DNA purified from Pf4 virions is not susceptible to trans-cleavage. (A) Plaque assay for LbCas12a PAM-dependent cis-cleavage. Bacterial lawns of *P. aeruginosa* PAO1 strain were induced to co-express LbCas12a and crRNAs complementary to Pf4 or JBD30 protospacers. Upon plating, bacteria were singly infected with serial dilutions of JBD30 or Pf4 phage. Black spots indicate bacterial clearance due to phage replication. (B) Plaque assay for ssDNA cleavage by LbCas12a. Bacterial lawns of *P. aeruginosa* PAO1 strain were induced to co-express LbCas12a and crRNAs complementary to Pf4 protospacers lacking a PAM. Upon plating, bacteria were singly infected with serial dilutions of JBD30 or Pf4 phage. crRNA complementary to a JBD30 protospacer with a PAM was used as a positive control for targeting. (C) Plaque assay for ssDNA trans-cleavage by LbCas12a. Bacterial lawns of *P. aeruginosa* PAO1 strain were induced to co-express LbCas12a and crRNA complementary to a JBD30 protospacer with a PAM. Upon plating, bacteria were singly infected or co-infected with JBD30 and Pf4 phage. Ten-fold serial dilutions of JBD were used while Pf4 was used at moderate and low titers. (D) *In vitro* LbCas12a cis- and trans-cleavage assay with dsDNA template (plus or minus protospacer) and DNA purified from Pf4 or M13 virions. (E) *In vitro* cleavage assay on DNA purified from PAO1 or Pf4 virions using enzymes as indicated. (F) *In vitro* cleavage assay on DNA purified from Pf4 virions using exonuclease I.

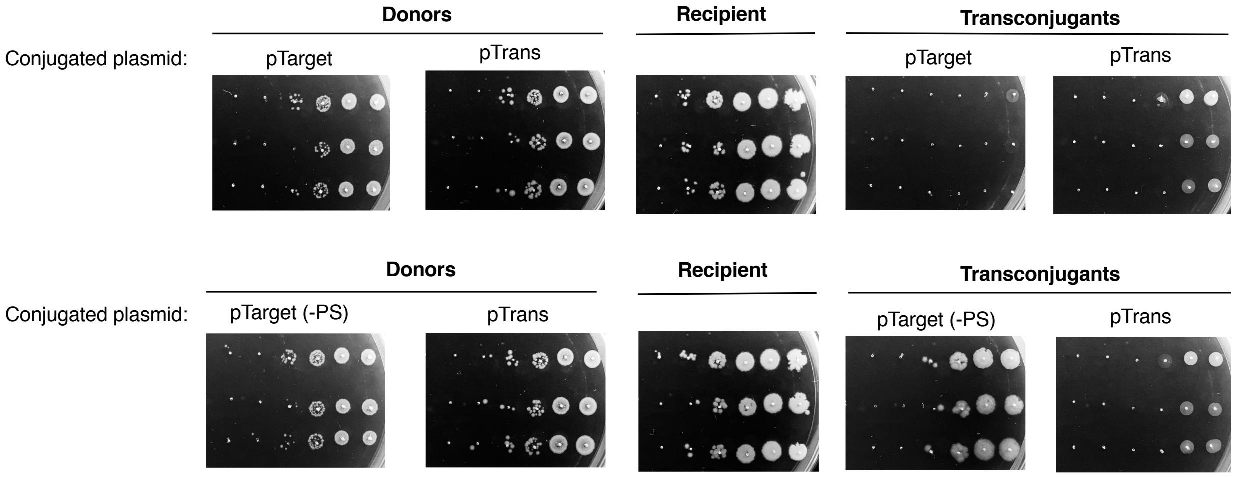


Figure S5. Conjugation assays for ssDNA trans-cleavage. The pictures show the colony forming units (10-fold dilution series) of donors (left), recipient (middle) and transconjugant (Right) cells grown in the appropriate selection media after the co-conjugation filter mating experiment (in triplicates: rows). The top panel shows the results for the co-conjugation of pTarget and pTrans. The bottom panel shows the results for the co-conjugation of pTarget(-PS) (same as pTarget but without the protospacer as a control for no targeting) and pTrans.

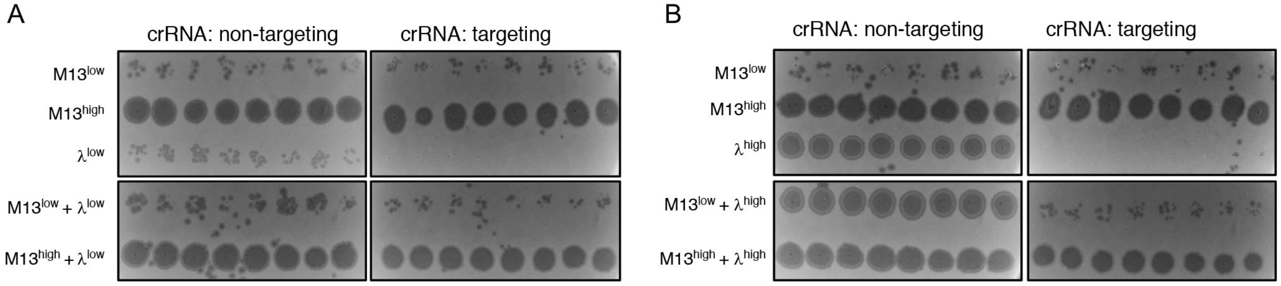


Figure S6. LbCas12a does not trans-cleavage M13 ssDNA during phage infection. Bacterial lawns of *E. coli* BW25113 F' strain were induced to co-express LbCas12a and non-targeting or λ-targeting crRNA. Upon plating, bacteria were singly infected or co-infected with low or high titers of M13 and (A) low or (B) high titers of $\lambda_{vir}$.

**Supplementary Table S1**

| **Strains** | **Species** |
| --- | --- |
| PAO1 | *Pseudomonas aeruginosa* |
| PAO1 tn7::LbCas12a | *Pseudomonas aeruginosa* |
| PAO1 tn7::LbCas12a attB::crRNA$_{DMS3}$ | *Pseudomonas aeruginosa* |
| BW25113 F' tn7::LbCas12a | *Escherichia coli* |
| S17-1 | *Escherichia coli* |
| CSH26 | *Escherichia coli* |

| **Phages** |
| --- |
| DMS3 |
| JBD30 |
| M13 |
| $\lambda_{vir}$ |
| Pf4 |

| **Plasmids** | | **Insert sequence** |
| --- | --- | --- |
| pHERD30T **(pTarget(-PS))** | | NA |
| pHERD20T | | NA |
| pKJK5 **(pTrans)** | | |
| *Insert: protospacer* | | |
| pHERD30T-ps$_{DMS3}$-PAM **(pTarget)** | | ttatgtgcggtatcgaaacgggc**gaaa** |
| pHERD30T-ps$_{DMS3}$ | | ttatgtgcggtatcgaaacgggc |
| pHERD30T-ps$_{DMS3}$-PAM-flip | | **tttc**gcccgtttcgataccgcacataa |
| pHERD30T-ps$_{DMS3}$-flip | | gcccgtttcgataccgcacataa |
| *Insert: crRNA* | **Target** | |
| pHERD30T-crRNA-DMS3 | DMS3/JBD30 | taatttctactaagtgtagatgcccgtttcgataccgcacataa |
| pHERD20T-crRNA-M13$_1$ | M13 | taatttctactaagtgtagatctgcaacagtgccacgctgagagc |
| pHERD20T-crRNA-M13$_2$ | M13 | taatttctactaagtgtagataacgaaccaccagcagaaga |
| pHERD20T-crRNA-λ | λ | taatttctactaagtgtagatgagggcagttgcggtcgtggaac |
| pHERD30T-crRNA-Pf4$_1$ | Pf4 | taatttctactaagtgtagatagacgttccggcgtcagcgcctg |
| pHERD30T-crRNA-Pf4$_2$ | Pf4 | taatttctactaagtgtagatgcaggctccagggagacagcgac |
| pHERD30T-crRNA-Pf4$_3$ | Pf4 | taatttctactaagtgtagatttttcatcatccacctcgcccat |
| pHERD30T-crRNA-Pf4$_4$ | Pf4 | taatttctactaagtgtagattcgggtcggaaggatcggtgttc |
| pHERD30T-crRNA-Pf4$_5$ | Pf4 | taatttctactaagtgtagatagaaggccagccccgccgcaatc |
| pHERD30T-crRNA-RFP | RFP | taatttctactaagtgtagatccgttacccaggactcctcc |

| **crRNA** | **Target** | **Sequence** |
| --- | --- | --- |
| crRNA-DMS3 | DMS3/JBD30 | uaauuucuacuaaguguagaugcccguuucgauaccgcacauaa |
| crRNA-M13$_1$ | M13 | uaauuucuacuaaguguagaucugcaacagugccacgcugagagc |
| crRNA-M13$_2$ | M13 | uaauuucuacuaaguguagauaacgaaccaccagcagaaga |
